# Visualization of seasonal phosphorus re-translocation, and expression profile of phosphate transporters in a shortened annual cycle system of the deciduous poplar tree

**DOI:** 10.1101/2020.04.12.038562

**Authors:** Yuko Kurita, Satomi Kanno, Ryohei Sugita, Atsushi Hirose, Miwa Ohnishi, Ayumi Tezuka, Ayumi Deguchi, Kimitsune Ishizaki, Hidehiro Fukaki, Kei’ichi Baba, Atsushi J. Nagano, Keitaro Tanoi, Tomoko M. Nakanishi, Tetsuro Mimura

**Author notes:** Author for correspondence: Yuko Kurita and Tetsuro Mimura.

## Abstract

Phosphorus (P) is an essential macronutrient for plant growth. In deciduous trees, P is remobilized from senescing leaves and stored in perennial tissues during winter for further growth. Annual internal recycling and accumulation of P is considered an important strategy to support vigorous growth of trees. However, the pathways of seasonal re-translocation of P and the molecular mechanisms of this transport have not been clarified. Here we show the seasonal P re-translocation route visualized using the real-time radioisotope imaging and the macro- and micro-autoradiography. We analyzed the seasonal re-translocation P in poplar (*Populus alba*. L) cultivated under “a shortened annual cycle system”, which mimicked seasonal phenology in a laboratory. From growing to senescing season, sink tissues of ^32^P and/or ^33^P shifted from young leaves and the apex to the lower stem and roots. The radioisotope P re-translocated from a leaf was stored in phloem and xylem parenchyma cells and redistributed to new shoots after dormancy. Seasonal expression profile of phosphate transporters (*PHT1*, *PHT5* and *PHO1* family) was obtained in the same system. Our results reveal the seasonal P re-translocation routes at the organ and tissue levels and provide a foothold for elucidating its molecular mechanisms.

## Introduction

Phosphorus (P) is an essential macronutrient for plant growth, however its availability in soil is quite limited (Schachtman *et al*., 1998). To utilize the valuable P, plants have various strategies for the uptake, translocation, storage and recycling of P. Re-translocation of P is a strategy for its internal recycling *in planta*. P is re-translocated from source tissues (e.g. old leaves) to sink tissues (e.g. young leaves, shoot apex) as phosphate (Biddulph *et al*., 1958b; Mimura *et al*., 1996; Himelblau & Amasino, 2001). It is supposed that such re-translocation of phosphate between source and sink occurs via the phloem. Both downward and upward P re-translocation via the phloem from leaves, and the lateral movement from the phloem to xylem have been demonstrated using microautoradiographic methods in an herbaceous plant, red kidney bean (Biddulph, 1956; Biddulph *et al*., 1958a). Although the molecular mechanisms of Phosphate (Pi) transport at the cellular level, and signaling of P status, are being elucidated through various studies in Arabidopsis and rice (Ham *et al*., 2018; Victor Roch *et al*., 2019; Wang *et al*., 2021), relatively less is known about lateral transport and re-translocation in tissues and organs (Aubry *et al*., 2019).

In deciduous woody plants, the internal recycling of various nutrients has been studied using radiolabeled materials. Carbon (C) and sulfur (S) were shown to be re-translocated from leaves in a leaf age-dependent manner in cottonwood and *Populus tremula* × *P. alba,* respectively (Dickson, 1989; Hartmann *et al*., 2000). These studies showed that expanding leaves and old mature leaves re-translocate nutrients mainly into sink organs in the upper apical parts and lower parts of the tree (lower stem and roots), respectively. Kavka & Polle showed that the absorption and allocation of ^33^P from roots depended on the P conditions to which the plant was exposed, while young leaves were usually the predominant aboveground sink (Kavka & Polle, 2016).

Besides this daily re-translocation, deciduous woody plants remobilize various nutrients from senescing leaves to perennial tissues during autumn for rescuing nutrients before the leaves fall (Chapin & Kedrowski, 1983; Aerts, 1996; Keskitalo *et al*., 2005; Netzer *et al*., 2017b,a). We previously reported that P is remobilized from leaves during autumn senescence in *Populus alba*, and stored as phytic acid (myo-inositol-1,2,3,4,5,6-hexakis phosphate: IP_6_) in protein storage vacuoles in phloem and xylem parenchyma cells in perennial tissues such as twigs during winter (Kurita *et al*., 2014, 2017). Phytic acid is well-known as a major P-storage form in seeds (Raboy *et al*., 2001). IP_6_ concentrations in the twigs decreased immediately in early spring, suggesting that IP_6_ is being degraded and P is being re-translocated for new growth. In last decade, Pi transporters (*PHT1-4*, *PHO*s) and their expression profiles under stress conditions and in different tissues have been characterized in poplar (Loth-Pereda *et al*., 2011; Kavka & Polle, 2016; Zhang *et al*., 2016). Loth-Pereda *et al*. showed that *PtPT1*, *PtPT5*, *PtPT6*, *PtPT9*, and *PtPT12* are upregulated in leaves during autumn senescence, suggesting that these transporters may play a role in P re-translocation from source to sink in *Populus trichocarpa* (Loth-Pereda *et al*., 2011). However, the seasonal P re-translocation routes and the molecular mechanisms that regulate the transport are still largely unknown.

In the present study, we demonstrated seasonal shifts in the re-translocation routes using radioisotope techniques and characterized the seasonal expression profile of phosphate transporters. Since radioisotopes cannot be used in the field, we used a shortened annual cycle system that mimics the seasonal phenology of trees in the laboratory (Kurita *et al*., 2014). This system consists of 3 stages. Stage 1 mimics spring/summer, stage 2 emulates autumn and stage 3 emulates winter, respectively (for detail, see method and Fig. S1c). After leaf fall in stage 3, plants were transferred to stage 1 again and bud burst induced. In this system, poplar trees showed normal seasonal events (leaf coloration, leaf fall, bud burst) and seasonal P recycling events (e.g., the P remobilization from senescing leaves, the storing of IP_6_ in dormant stems and the redistribution during budburst)(Kurita *et al*., 2017). By using this system, the real-time behavior of P remobilized from a leaf, seasonal changes of re-translocation routes along the longitudinal and radial axis of stem, and overwintering recycling were successfully visualized. Further, we have analyzed changes in gene expression of leaves and stems under a shortened annual cycle system, focusing on Pi transport mechanisms. We discuss a relationship between P re-translocation visualized by radionuclides and molecular mechanisms suggested by gene expression analysis.

## Materials and Methods

### Plant materials and culture conditions

Poplar shoots with 4-5 leaves were cut and rooted in 1/5 MS medium, and transferred into the pots containing vermiculite, that had been washed with tap water ten times and by distilled water five times. Potted cuttings were placed in tree culture chambers (LH-350S, NK system, Osaka, Japan) in turn from stage 1 to 3 under the conditions in ‘a shortened annual cycle system’ as described before (Kurita *et al*., 2014)(Fig. S1c). Briefly, stage 1 (25℃, 14h light) mimics spring/summer condition, and promotes vigorous plant growth. Stage 2 (15℃, 8h light) mimics autumn condition, and causes plants to experience short daylength and moderate cooler temperatures. Stage 3 (5℃, 8h light) mimics winter condition, and promotes leaf senescence. At this stage, plants cease apical growth. The leaves turn yellow and then fall.

Plants were fertilized with 1/5 MS medium (1L / 6 potted cuttings) once a week at stage 1. Afterwards they were cultured with distilled water only. In stages 1 and 2, plants at 3^rd^ to 4^th^ week, and in stage 3, plants at 3^rd^ to 5^th^ week, were used for the experiments.

### Imaging using the real-time radioisotope imaging system

To add radiophosphorus, a ‘flap’ was cut along a vein of 6^th^ leaf in the manner shown in Fig.S1a (Biddulph & Markle, 1944), and 5 μl ^32^P-phospate (H_3_ ^32^PO_4_) solution in 0.2M KH_2_PO_4_ (final concentration of 200 kBq μl^-1^) was added. After 1 or 3 min, the flap was cut off because its strong signals disturb the imaging. Then, changes in ^32^P distribution were observed using the real-time radioisotope imaging system established previously (Kanno *et al*., 2012; Hirose *et al*., 2013; Sugita *et al*., 2013, 2016). The plant was mounted on the fiber optic plate coated with 100 μm thickness of CsI (Tl) scintillator by vapor deposition (FOS, 100 mm × 100 mm × 5 mm, Hamamatsu Photonics Co., Hamamatsu, Japan). Two fiber optic plates were connected longitudinally. The beta ray emitted from ^32^P in the sample interacts with the CsI (Tl) scintillator, and then is converted to visual light. Light was captured by the cooled CCD camera with an image capture system (Aquacosmos/VIM system; resolution: 640 pixel × 480 pixel, Hamamatsu Photonics Co., Hamamatsu City, Japan) (Kanno *et al*., 2012; Sugita *et al*., 2013). The measurement was performed for three hours in a dark room at 20°C. The same measurements were repeated 3 times.

To analyze the velocity of ^32^P transported in the vascular bundle, the signal intensity of the region of interest (ROI) was measured by ImageJ software. Two ROIs were set at the apex and the 6^th^ leaf axil. Three ROIs were set at corners of FOS to measure background signals. Of the 3 replicates, the 2 trees which showed the highest signals were analyzed. The velocity was calculated from the distance between the apex and the 6^th^ leaf axil and the difference between the times it took for the signal to reach the detection threshold. The detection threshold was defined as five times the standard deviation of the background signal intensity.

### Whole plant autoradiography using imaging plate

^32^P-phospate (5 μl) in 0.2M KH_2_PO_4_ (final concentration of 200 Bq lμ^-1^) was added to 6^th^ leaf as described above. Then, plants were cultured in each stage condition for 1 day. After 1 day culture, the 6^th^ leaf was cut off because its strong signals disturb the imaging. The vermiculite was washed away from poplar roots with tap water. Plants were then mounted on commercial xerographic paper, and wrapped with double layers of polyvinyl film. The sample was set on an imaging plate (Fujifilm, Tokyo, Japan) in the cassette case and exposed in a dark room for 3 days. The imaging plate was then measured using Typhoon FLA 9500 (GE Healthcare UK Ltd., Buckinghamshire, England). The same measurements were repeated 3 times.

### Whole plant autoradiography using imaging plate with phloem girdling

Approximately 2 mm of phloem from the upper or lower internodes of the 6^th^ leaf of a plant at stage 1 was cut with a razor blade and peeled off, effectively disrupting phloem translocation in this region (Fig. S1b). Then immediately, 5 μl ^32^P-phospate (KH_2_ ^32^PO_4_) solution in 0.2 M KH_2_PO_4_ (final concentration of 1 kBq μl^-1^) was added to the 6^th^ leaf as described above. Plants were cultured in stage 1 for 3 h or 1 day. After culture, the 6^th^ leaf was cut off. Upper and lower internodes of 6^th^ leaf were cut to prevent re-translocation of ^32^P during contact. The vermiculite was washed away from poplar roots with tap water. Plants were then mounted on commercial xerographic paper and wrapped with double layers of polyvinyl film. The sample was set on the imaging plate in the cassette case and exposed in a dark room for 1 day. Then the imaging plate was measured using Typhoon FLA 9500. The same measurements were repeated 3 times.

### Microautoradiography (MAR) of the stem

^33^P-phospate (5 μl) in 0.2M KH_2_PO_4_ (final concentration of 37 or 74 kBq μl^-1^) was added to 6^th^ leaf as described above. Then, plants were cultured in each stage condition for 1 day. ^32^P is not suitable for obtaining high-resolution images because its energy is too high. Therefore, the experiments were performed by using ^33^P which has a lower energy. ^33^P distribution in stems was detected by the MAR method for fresh-frozen sections, as described previously (Hirose *et al*., 2014). Stem samples that were cut from a poplar in each stage were frozen with liquid nitrogen. The frozen stems were embedded in a dedicated embedding medium (SCEM; SECTION-LAB Co. Ltd.) and sliced into 20 μm thick sections using a Cryostat (CM1850; Leica Microsystems, Wetzlar, Germany) set at −20□. Sectioning was performed using the Cryo-Film transfer kit (SECTION-LAB Co. Ltd., Hiroshima, Japan) (Kawamoto, 2003; Hirose *et al*., 2014).

Frozen sections were put on hydrophilic amino-coated glass slides (MAS-GP type A; Matsunami Glass Ind., Ltd., Osaka, Japan) coated with a photosensitive nuclear emulsion (Ilford Nuclear Emulsion Type K5; Harman Technology Ltd., Cheshire, UK) in the Cryostat, and contacted tightly at −80□ for approximately 1 month or more. A 1.2 μm thick polyphenylene sulfide (PPS) film was inserted between the frozen sample sections and the photosensitive nuclear emulsion on the glass slide to enable easy separation of the section and the emulsion after exposure. The results scarcely changed when the sample was exposed for more than 1 month.

After the exposure, the sections were removed from the glass slides in the cryostat set at −20□ in the darkroom. The glass slides were soaked in quarter-strength D-19 developer (Eastman Kodak Co., NY, USA) for 6 min at room temperature to develop the silver particles. Then, the slides were soaked in distilled water for 1 min to stop the development, followed by exposure to 0.83 M sodium thiosulfate for 4 min for fixation. Finally, the glass slides were soaked in distilled water for 10 min twice. The temperature of the darkroom was maintained at 20□ to ensure development.

In order to stain the sliced tissue, the sections were floated on distilled water for 1 min three times at room temperature to wash out the embedding material. Then the sections were stained with 0.25 % toluidine blue for 30 sec and rinsed with water for 1 min. The sections were then mounted on a new glass slide using a glycerol-based mounting medium (SCMM; SECTION-LAB Co. Ltd.).

The stained sections and the silver particles developed by MAR were observed and photographed using a microscope (BX-60; Olympus, Tokyo, Japan). The tissue images and radiographs were then assembled using e-Tiling (Mitani Co., Tokyo, Japan) or ImageJ (http://rsb.info.nih.gov/ij) by its plugin, Grid/Collection stitching software (Preibisch *et al*., 2009). The registration was performed on the tissue images and radiographs using ImageJ and LPixel’s ImageJ Plugin, Lpx Registration software (https://lpixel.net/products/lpixel-imagej-plugins/) and GIMP software (http://www.gimp.org). The same measurements were repeated 3 times.

### Serial autoradiography of the leaf axil and 3D reconstruction

Fresh-frozen sections from the axil of the 6^th^ leaf were prepared for autoradiography experiments using the imaging plate (BAS IP MS; GE Healthcare UK Ltd.). The thickness of the section was 20 μm. The exposure time was about 76 h for samples in stage 1 at −80□. In stage 3, exposure time was about 22 h because the signal was stronger than that of samples in stage 1. The imaging plate was scanned with an FLA5000 image analyser (Fujifilm, Tokyo, Japan) at a resolution of 10 µm.

After linear contrast enhancement and registration using ImageJ pluguin StackReg (Thévenaz *et al*., 1998), autoradiographs were converted to 3D images using ImageJ plugin Volume Viewer (http://rsb.info.nih.gov/ij/plugins/volume-viewer.html) and 3D Viewer (http://imagej.nih.gov/ij/plugins/3d-viewer/). The same measurements were repeated 3 times.

### Whole plant autoradiography using imaging plate over dormancy period

^32^P-phospate (7 μl) in 0.2M KH_2_PO_4_ (final concentration of 700k Bq μl^-1^) was added to the 6^th^ leaf of a plant at the 4^th^ week in stage 3 as described above. After leaf fall (at 8^th^ week in stage 3) or after bud burst (at 2^nd^ week in stage 1 of 2^nd^ cycle), the vermiculite was washed away from poplar roots with tap water. The plant was then mounted on paper, and wrapped with a double layer of polyvinyl film. The sample was set on the imaging plate in the cassette case and exposed in a dark room for 15 min or 1 day. Then, the imaging plate was measured using Typhoon FLA 9500. The same measurements were repeated 3 times.

### RNA-Seq analysis

The 6^th^ leaf and stem (4-8 cm from apex) without node were sampled between 13:00 and 15:00 (stage 1: 6-8 hours after lighting, stages 2 and 3: 3-5 hours after lighting), immediately frozen in liquid nitrogen, and stored at −80 °C. At stage 1 in the 2^nd^ cycle, the stem at 4-8 cm from the apex of the end of the 1^st^ round were sampled. A summary of sample sets is shown Fig. S5a. Leaf samples were ground with a mortar and pestle in liquid nitrogen. Stem samples were ground using the Multi-Bead Shocker (MB601KU, Yasui Kikai Co.). Total RNA was extracted using the RNeasy Plant Mini Kit (Qiagen, USA). For RNA-Seq library preparation, 500 or 1000 ng of total RNA per sample was used. We prepared the library according to the procedure described previously (Nagano *et al*., 2015). The quality of the library was assessed by the Bioanalyzer (Agilent Technologies, USA). The libraries were sequenced by the single-read 50 bases with the HiSeq 2500 (Illumina, USA).

Preprocessing and quality filtering of RNA-Seq data were performed using the Trimmomatic-0.33 (Bolger *et al*., 2014). Then, preprocessed reads were mapped on primary transcript sequences in *Populus trichocarpa* v3.1 (downloaded from Phytozome v12.1) (Tuskan *et al*., 2006) with Bowtie1(v1.1.1) (Langmead *et al*., 2009) and quantified using RSEM-1.2.21(Li & Dewey, 2011). The output of RSEM was analyzed using R software (R Core Team, 2019).

Seasonal expression analysis was performed on each of the 6^th^ leaf and the stem data as follows. Samples with fewer than 10^6^ reads were excluded from the analysis. Attributes of the samples are shown in Supplementary table S1. The analysis was then conducted with reference to the pipeline proposed by Law et al.(Law *et al*., 2018). The expressed genes (the 6^th^ leaf: 23646 genes, Stem: 25151 genes) were defined using filterByExpr function of the edgeR package (Robinson *et al*., 2010; Chen *et al*., 2016) that excluded genes that did not have sufficient read counts.

After removing heteroscedascity from count data using the voom function of the limma package (Law *et al*., 2014), seasonality-induced genes (SI genes) which show a quasi-seasonal oscillation in gene expression in shortened annual cycle system were detected by an ANOVA-like test in limma (thresholds of topTableF function: p.value = 0.01, lfc = 1)(Ritchie *et al*., 2015). In the 6^th^ leaf and stem, 15743 and 17234 genes were detected as SI genes, respectively.

After dividing the expression patterns of SI genes using k-means clustering, GO enrichment analysis was performed on each cluster. The GO annotations of *Arabidopsis thaliana* ortholog obtained from TAIR10 was used for the analysis. *Arabidopsis thaliana* orthologs were obtained from ‘best Athaliana TAIR10 hit name’ of Ptrichocarpa_444_v3.1 annotation_info file. Statistical tests of enrichment analysis were performed by the Fisher’s exact test function provided by the R software. Multiple testing correction was performed by FDR (Benjamini & Hochberg, 1995). GO terms with adjusted *P* < 0.05 are defined as enriched GO terms. Representative GO terms were summarized using REVIGO (http://revigo.irb.hr) (Supek *et al*., 2011) (allowed similarity = 0.5) and visualized with reference to the previous protocol (Bonnot *et al*., 2019).

For *PHT* and *PHO* family genes detected as SI genes, Tukey HSD were performed in R to test for differences between sampling points (*P* < 0.05).

## Results

### Shift of longitudinal re-translocation routes of phosphorus between growing and senescence seasons

To continuously observe how P is re-translocated from a leaf, we performed real-time radioisotope imaging using plants grown in the shortened annual cycle system. In stage 1, ^32^P was re-translocated from the 6^th^ leaf in both upward and downward directions, and gradually accumulated at apical parts (apex and young leaves) in 3h (Fig.1a, Video S1, Fig. S2). Within 30 min, ^32^P was detected at the leaf axil of the applied leaves, and after 1 to 1.5 h, ^32^P was detected at the shoot apex. Estimated transport velocities were about 0.015 mm s^-1^ in tree A and 0.021 mm s^-1^ in tree B (Fig. S2). On the other hand, in stage 3, ^32^P was only re-translocated downwards from the 6^th^ leaf. No signals in upper parts were detected within 3h (Fig.1b, Video S2). To observe ^32^P transport and distribution in whole plants over longer periods, plants were cultured for 1 day after ^32^P application, and contacted to imaging plates (Fig.2). In stage 1, ^32^P was detected mainly in the upper parts, in agreement with the results of the real-time imaging experiment. These results correspond with previous studies on re-translocation of C and S from leaves (Dickson, 1989; Hartmann *et al*., 2000). During stages 2 and 3, ^32^P was detected mainly in the lower parts (stem and roots), and slightly in the upper stem.

**Fig. 1.**
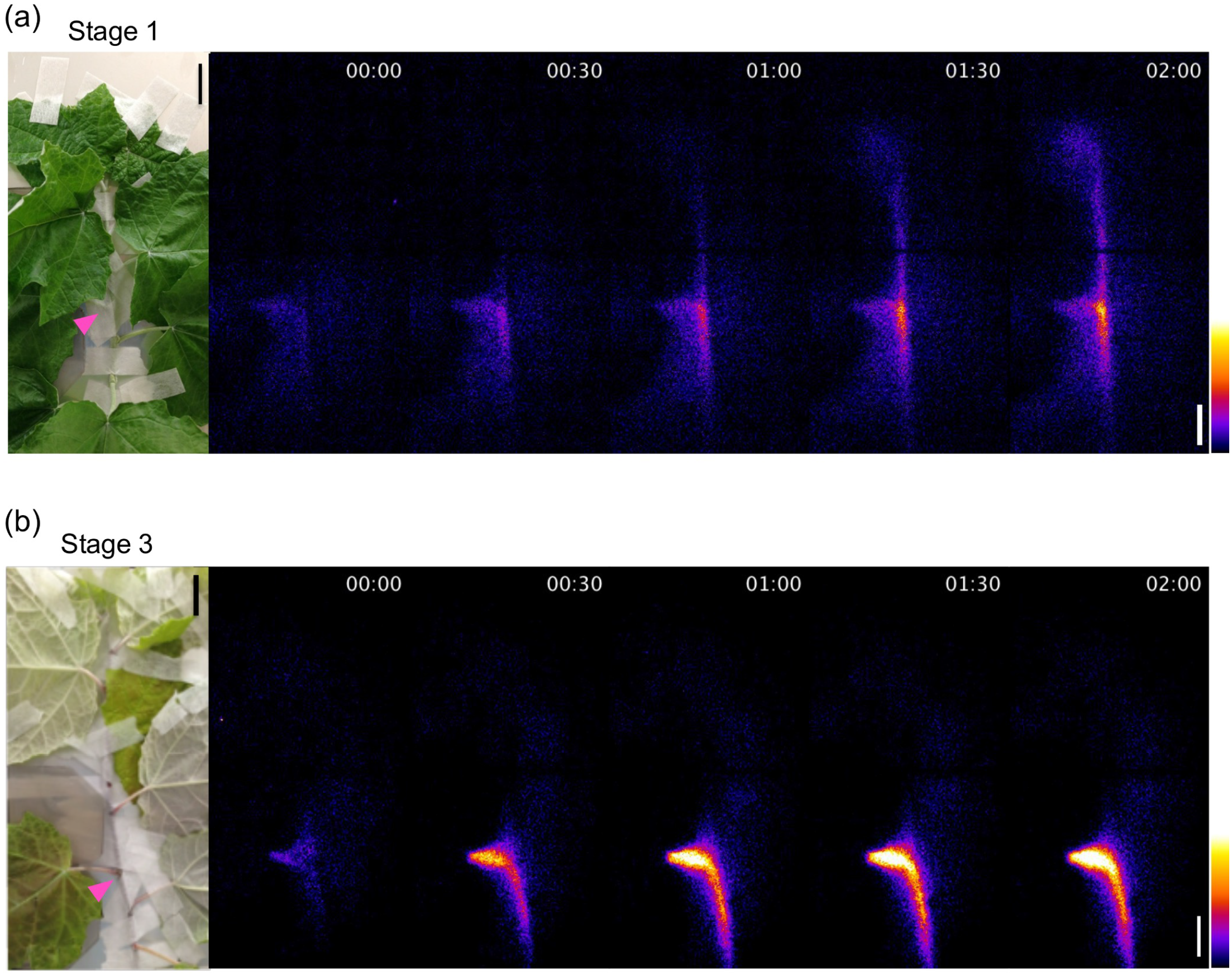
Real-time imaging of ^32^P in *Populus alba*. (a) At stage 1. (b) At stage 3. Pink color arrowheads indicate the position of the 6^th^ leaf. The 6^th^ leaf was shielded by acrylic board because its strong signals disturbed measurement. bar = 2 cm.

**Fig. 2.**
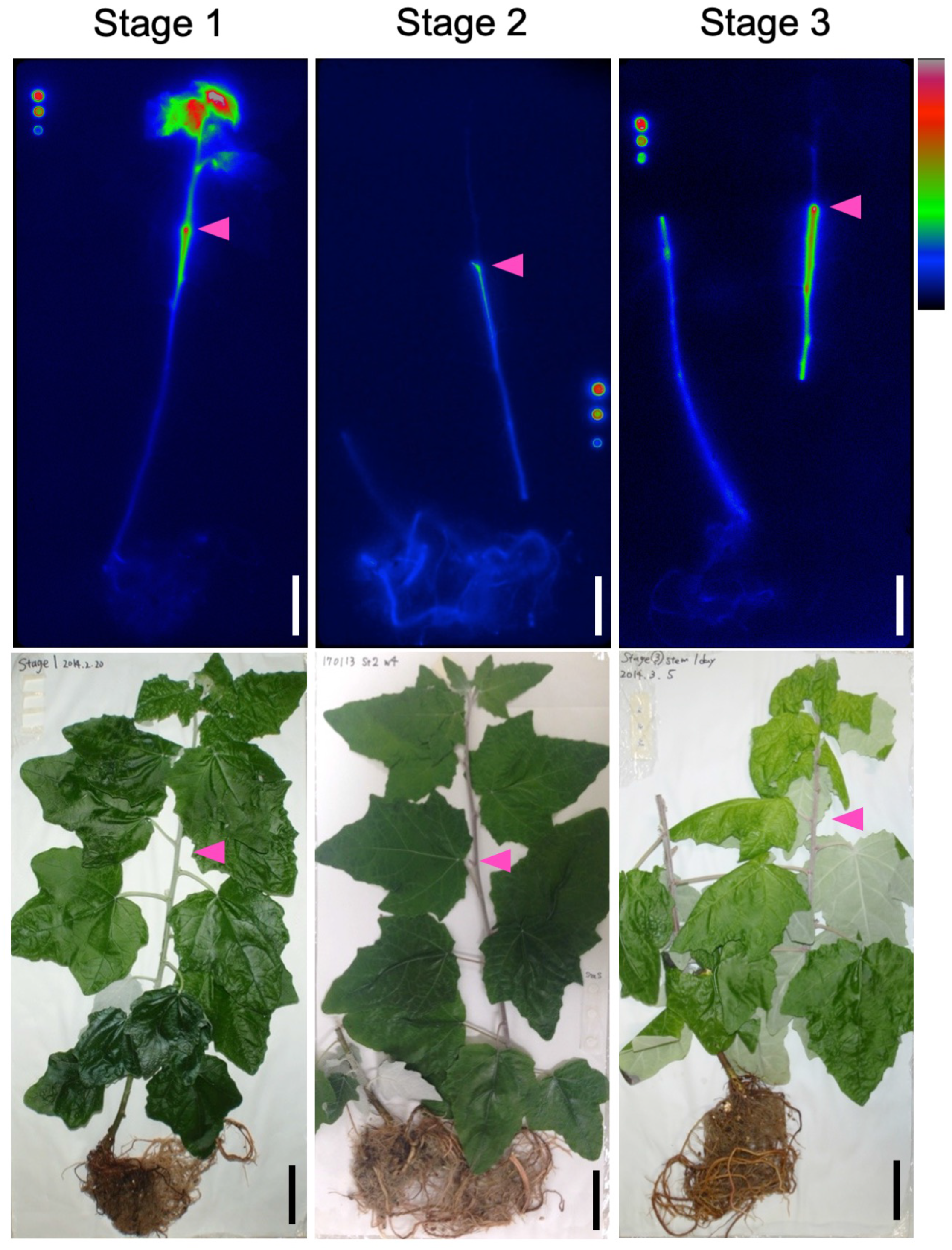
Whole plant autoradiographs of *Populus alba* trees in a shortened annual cycle system. Images after one day of ^32^P application at each stage. Pink color arrowheads indicate the position of the 6^th^ leaf. The 6^th^ leaf was cut off because its strong signals disturbed measurement. In stage 2 and 3, the stem was cut because the plant was too large for the imaging plate. bar = 4 cm.

### Longitudinal re-translocation of phosphorus via phloem

To reveal which tissue (xylem and/or phloem) is involved in longitudinal re-translocation of P, changes in the re-translocation route of P were observed by phloem girdling using a tree in stage 1 (Fig. 3, S1b). The upper or lower phloem of a node attached to the 6^th^ leaf was peeled off, and then ^32^P was immediately applied to the 6^th^ leaf. After 3 h or 1 day culture, the plant was set on imaging plate for ^32^P detection. When the phloem was peeled off above the applied leaf, ^32^P was detected only in the lower stem and roots after 3 h (Fig.3a). After 1 day, ^32^P was detected mainly in the lower stem and roots, and slightly in the upper stem and young leaves. When phloem was peeled off below the applied leaf, ^32^P was detected only in the upper stem and young leaves after 3 h or 1 day culture (Fig.3b). These results suggest dominant roles of phloem in longitudinal re-translocation of P in *Populus alba*.

**Fig. 3.**
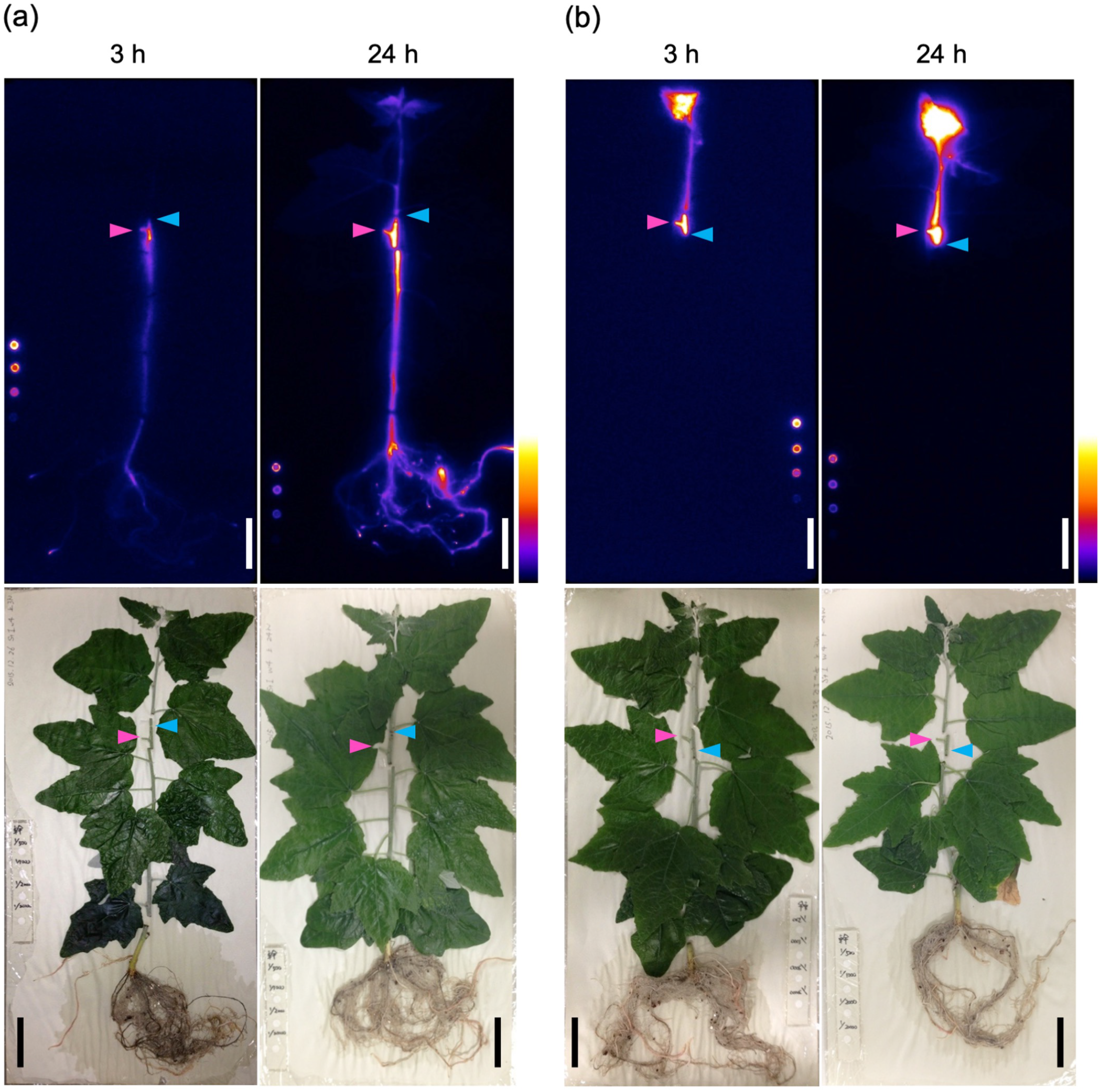
Whole plant autoradiographs of *Populus alba* trees with phloem girdling. **(a)** Plants peeled of their upper phloem from the 6^th^ leaf were cultured for 3 h or 24 h. **(b)** Plants peeled of their lower phloem from the 6^th^ leaf were cultured for 3 h or 24 h. Plants at stage 1 were used for experiments. ^32^P solution was added to the 6^th^ leaf indicated by pink color arrowheads. Blue color arrowheads indicate the position of phloem girdling. After cultivation, upper and lower internodes of the 6^th^ leaf were cut to prevent further re-translocation of ^32^P during contact. bar = 4 cm.

### Visualization of P inflow from a leaf to the stem

To observe P inflow from a leaf to the stem in detail, serial autoradiographs of a leaf axil were converted to 3D images (Fig. 4a, b and S3). ^33^P was used for more detailed observation in these experiments. In each stage, plants were cultured for 1 day after ^33^P application. In stage 1, ^33^P was detected throughout the stem, petiole and bud. In the stem, ^33^P was mainly distributed in the phloem region on the side of applied leaf (Fig. 4a, S3a, Video S3). On the other hand, in stage 3, ^33^P was detected intensively in the lower phloem on the applied side of the leaf and petiole (Fig. 4b, S3b, Video S4). ^33^P was barely detected in the upper stem, axillary bud or the phloem opposite the petiole. The upward transport clearly ceased at stage 3 compared with stage 1. At the junction between the petiole and the stem, there appeared to be formation of an abscission layer (Fig. 4b, S3b).

**Fig. 4.**
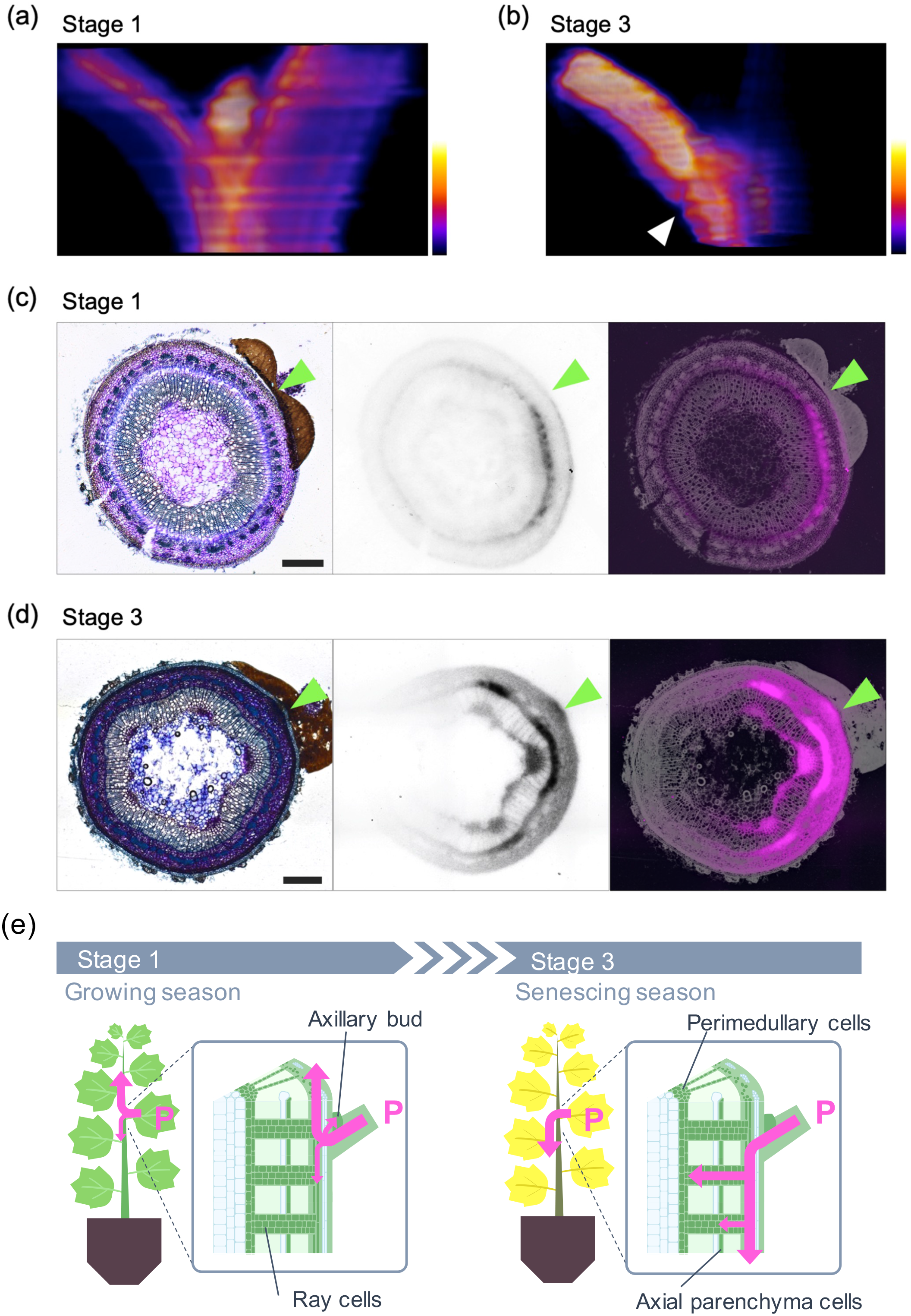
Side view of the 6^th^ axil and microautoradiographs of the stem between 6^th^ and 7^th^ nodes of *Populus alba*. (a) Side view of the 6^th^ axil at stage 1. ^33^P autoradiograph sections were converted to 3D images using ImageJ. (b) Side view of the 6^th^ axil at stage 3. The white color arrowhead indicates abscission layer during formation. (c) The stem between 6^th^ and 7^th^ nodes in stage 1. Left: the section stained by toluidine blue. Middle: Microautoradiograph of the section. Right: merged image. Green color arrowheads indicate the position of the 6th leaf. (d) The stem between 6^th^ and 7^th^ nodes in stage 3. Bar = 0.5 mm. (e) A schematic diagram of the seasonal change of P re-translocation in *Populus alba*. In the growing season, P is mainly re-translocated from the 6^th^ leaf to upper tissues (young leaves, apex) via phloem. P is re-translocated also to the axillary bud. During senescence, P is re-translocated from the 6^th^ leaf to lower tissues (stem, roots) via phloem. P is exchanged from phloem to xylem near the 6^th^ leaf and accumulated in ray cells and perimedullary cells.

### Shift of radial P re-translocation from a leaf between growing and senescence seasons

In the stem of woody plants, cells form three-dimensional lattices and they transport water and solutes along both the longitudinal axis and radial axes (Van Bel, 1990; Spicer, 2014). To observe radial transport inside a stem during senescence, we used microautoradiography because of its higher resolution compared to conventional autoradiography using an imaging plate. Plants on which ^33^P was applied to the 6^th^ leaf were cultured for one day. ^33^P localization in fresh frozen sections of the stem was detected. In the growing season at stage 1, ^33^P was detected mainly in phloem and to a lesser extent throughout the section of the stem just below the 6^th^ leaf (stem between 6^th^ and 7^th^ nodes) (Fig. 4c). As shown in the 3D reconstruction (Fig. 4a), the highest signals were recorded in the phloem on the side of the ^33^P applied leaf. This tendency was also observed in the upper stem (stem between 5^th^ and 6^th^ nodes) and the lower stem (stem between 9^th^ and 10^th^ nodes) (Fig. S4a, b).

During senescence at stage 3, ^33^P was strongly detected in the phloem on the side of the applied leaf, and also in ray tissues and parenchyma cells around the pith (perimedullary cells) (Fig. 4d). ^33^P was slightly detected both in the upper stem (stem between 5^th^ and 6^th^ nodes) and the lower stem (stem between 9^th^ and 10^th^ nodes) (Fig. S4c, d). Because ^33^P accumulation in xylem parenchyma cells was stronger in the stem between 6^th^ and 7^th^ nodes than in the stem between 9^th^ and 10^th^ nodes, the exchange from phloem to xylem probably occurred in the stem nearby the applied leaf rather than in the lower stem and roots (Fig. 4e).

### Re-utilization of overwintering P stored in stem and roots by new growing tissues in spring

To reveal how P stored in winter is re-utilized in spring, artificial overwinter experiments were performed in the shortened annual cycle system (Fig. 5). Plants, which had ^32^P applied to the 6^th^ leaf were cultured for about 1 month until leaf fall in stage 3. Just after leaf fall, ^32^P was detected mainly in the lower stem and roots (Fig. 5a). Other trees were moved to stage 1 and bud burst induced. After bud burst in plants put in the second shortened annual cycle, ^32^P remobilized from the 6^th^ leaf was detected throughout the whole plant, particularly at apices and young leaves of new shoots (Fig. 5b).

**Fig. 5.**
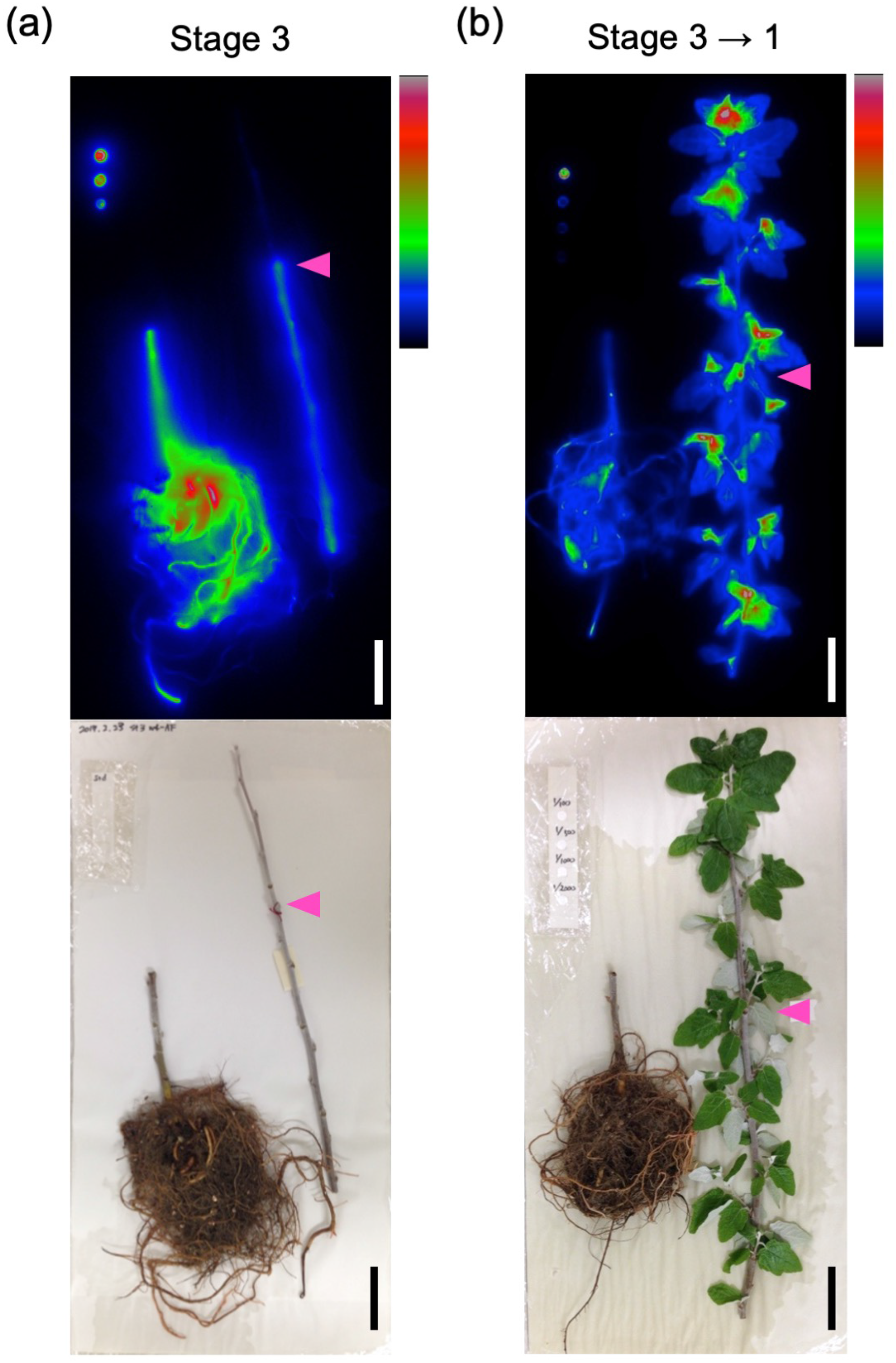
Whole plant autoradiographs of *Populus alba* trees over dormancy period in the shortened annual cycle system. Poplar trees in which ^32^P was applied to the 6^th^ leaf at 4^th^ week in stage 3 were subsequently cultured in stage 3. (a) Image after 4 weeks of ^32^P application (just after leaf fall). (b) Image after 6 weeks of ^32^P application. After 4 weeks cultivation, plants were moved to stage 1, and bud burst was induced. Pink color arrowheads indicate the position of the 6^th^ leaf. The stem was cut because the plant was too large for the imaging plate. bar = 4 cm.

### Quasi-seasonal expression patterns in the shortened annual cycle system

In order to investigate molecular system of P re-translocation, RNA-Seq analysis was performed to investigate the time course of gene expression in the 6^th^ leaf and the stem of poplar grown in the shortened annual cycle system. In the 6^th^ leaf and stem, 15,743 and 17,234 genes were detected as seasonality-induced genes (SI genes) which shows a quasi-seasonal oscillation in gene expression in shortened annual cycle system (Supplementary table S2-3). We classified SI genes in the 6^th^ leaf and stem into three and five clusters, respectively, using the k-means clustering method (Fig. 6ab). Then, GO enrichment analysis was performed for each cluster, and enriched biological process GO terms were summarized (Fig. 6cd, Supplementary Table S4-11).

In the 6^th^ leaf, SI genes that showed higher expression during growing season were classified into cluster A (Fig. 6a). In this cluster, enrichments of GO terms such as ‘organonitrogen compound biosynthetic process’ (GO:1901566), ‘photosynthesis’ (GO:0015979) and ‘cell wall organization or biogenesis’ (GO:0071554) were enriched (adjusted *P* value < 0.05, See Supplementary Table S4-6 for exact values.) (Fig. 6c). SI genes that showed higher expression during the senescing season were classified into cluster C. GO terms such as ‘autophagy’ (GO:0006914), ‘catabolic process’ (GO:0009056) and ‘transport’ (GO:0006810) were enriched in this cluster, and suggested that leaf senescence occurred at stage 3. GO terms such as ‘phosphorus metabolic process’ (GO:0006793), ‘cellular response to phosphate starvation’ (GO:0016036) and ‘phosphate ion homeostasis’ (GO:0055062) were also enriched. ‘cellular response to phosphate starvation’ and ‘phosphate ion homeostasis’ were represented by “response to extracellular stimulus” (GO:0031668) and ‘anion homeostasis’ (GO:0055081) (Supplementary Table S6).

**Fig. 6.**
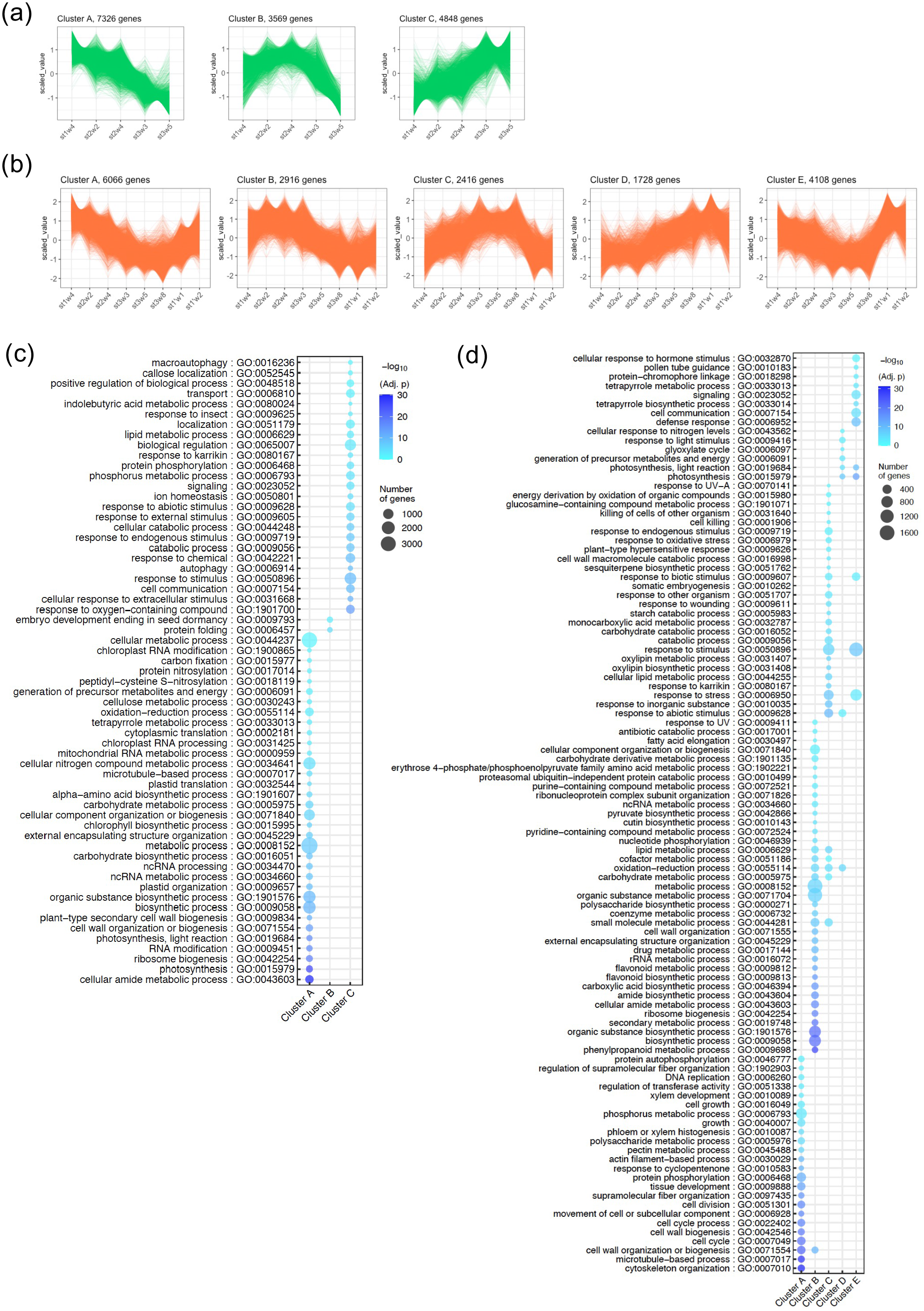
Clustering plots and GO enrichment analysis of RNA-Seq data. (a) Clustering plots of seasonality-induced genes (SI genes) in the 6^th^ leaf. (b) Clustering plots of SI genes in stem. (c) Enriched biological process gene ontology (GO) terms in SI genes in the 6^th^ leaf (adjusted *P* < 0.05). (d) Enriched biological process GO terms in SI genes in stem (adjusted *P* < 0.05). The circle size indicates the number of genes. Color scale indicates −log_10_ (adjusted *P*) in GO enrichment analysis.

About stem samples, we added sampling points at 8^th^ week in stage 3 and during stage 1 in second cycle (st1’w1 and st1’w2) to leaf samples. The clusters are shown in order from A to E according to the position of the peaks in the expression pattern (Fig. 6b). In cluster A, GO terms suggesting active growth such as “cell wall biogenesis” (GO:0042546), “cell division” (GO:0051301) and “growth” (GO:0040007) were enriched (adjusted *P* value < 0.05, See Supplementary Table S7-11 for exact values.). “cell wall organization or biogenesis” (GO:0071554) was commonly detected in clusters A and B. In cluster C, GO terms such as “response to abiotic stimulus” (GO:0009628) and “response to cold” (GO:0009409) represented by “response to Karrikin” (GO:0080167) were enriched (Supplementary table S9). Previous studies have shown that the expression of defense-related genes is up-regulated in early winter and that the expression of signaling-related genes is up-regulated during late winter and early spring in bark of *Populus deltoides* (Park *et al*., 2008). Enrichment of GO terms such as “killing of cells of other organism” (GO:0031640) and “immune response” (GO:0006955) (represented by “plant-type hypersensitive response” (GO:0009626)) in cluster C, “signaling” (GO:0023052) and “cell communication” (GO:0007154) in cluster E were consistent with that. “defense response” (GO:0006952) was also enriched in cluster E. GO terms such as “photosynthesis” (GO:0015979) were commonly enriched in Cluster D and Cluster E. This may suggest that photosynthesis in the stem of this system play a role in energy metabolism during the leafless period.

We confirmed the expression pattern in our laboratory system for genes whose seasonal expression patterns have been investigated in the field in previous studies (Fig. 7a, S7b). Lu *et al*. reported that the expression of *LHB1B1* and *CA1* decreased from July to Oct, while the expression of *GLR2* and *WRKY* increased in *Populus trichocarpa* Nisqually-1 by RT-qPCR and RNA-Seq (Lu *et al*., 2020). Seasonal expressions of *DPD1* in a shortened annual cycle and in field has been investigated previously by qPCR, and shown to increase before leaf fall (Takami *et al*., 2018). In our study, *LHB1B1* and *CA1* showed the highest gene expression at stage 1, and were classified as cluster A. *WRKY75* and *DPD1* showed highest gene expression at stage 3, and were classified as cluster C. Although *GLR2* was expressed, the variability of its expression was large, and was not identified as a SI gene.

**Fig. 7.**
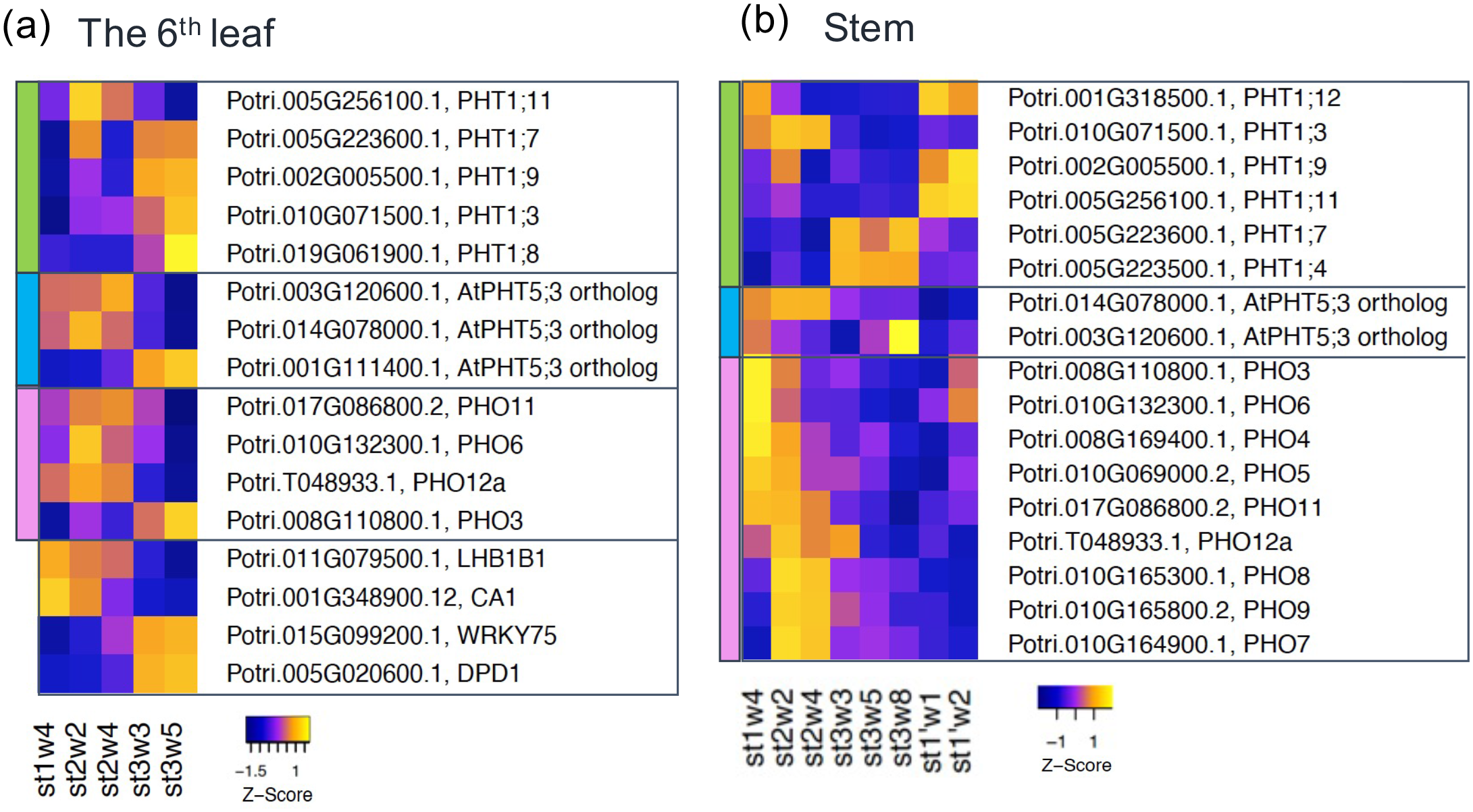
Gene expression of phosphate transporters in *Populus alba*. (a) Expression heatmap of phosphate transporters and typical seasonally variable genes in the 6^th^ leaf. (b) Expression heatmap of phosphate transporters in the stem. The color of the leftmost bar indicates the group of genes. Green: *PHT1*, Blue: *PHT5*, Pink: *PHO*. The mean value of log_2_(cpm) (prior.count = 2) was normalized by the scale function so that the mean is 0 and the variance is 1. See Fig. S6-9 for plots of the expression levels of individual genes.

### Expression patterns of phosphate transporters in the 6^th^ leaf

To investigate the molecular mechanisms underlying the seasonal shift of P re-translocation route, expressions of phosphate transporters (*PHT1*, *PHT5*, and *PHO*) were examined (Fig. 7ab, S6-9, Table 1). The gene numbers of *PtPHT1* family and *PtPHO* family (homologs of *AtPHO1* family) refer to the previous studies by Kavka & Polle and Zhang *et al*., respectively (Kavka & Polle, 2016; Zhang *et al*., 2016). *PHT1*s are mainly localized in the plasma membrane for the uptake of Pi into the cell, while *PHT5*s are mainly localized in the vacuole (Liu *et al*., 2015, 2016; Wang *et al*., 2021). *PHT5*s themselves have not previously been identified and analyzed in *Populus*. Therefore, we selected genes which were annotated as the ortholog of *AtPHT5;1-3* based on ‘best Athaliana TAIR10 hit name’ in the Phytozome database. There were 3 transcripts that were described that ‘best Athaliana TAIR10 hit name’ is *PHT5;3* (Potri.001G111400.1, Potri.003G120600.1 and Potri.014G078000.1). No genes were annotated as orthologs of AtPHT5;1 or AtPHT5;2.

**Table 1.**
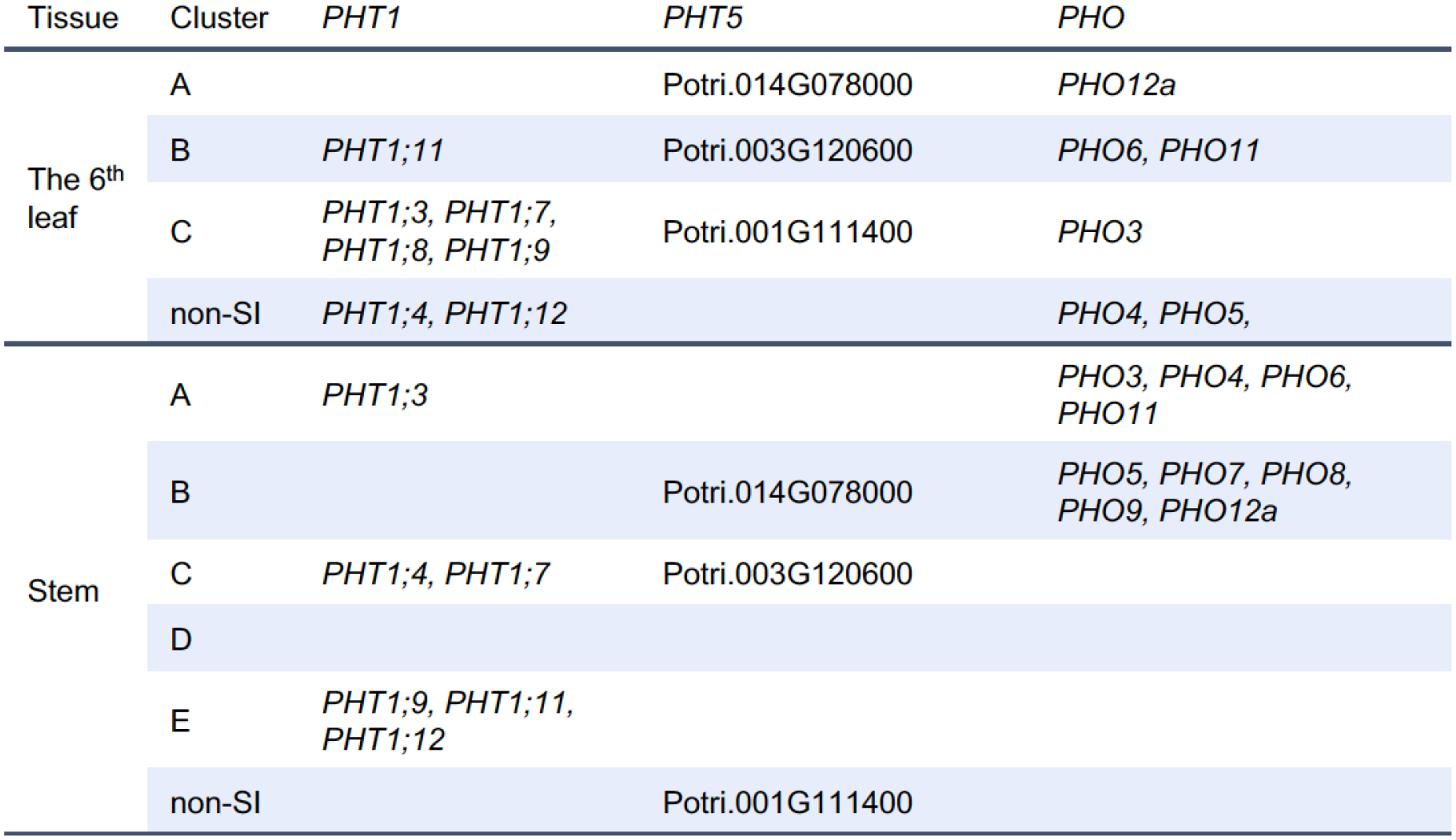
Phosphate transporters classified into clusters in Fig. 6. “non-SI” indicates genes that showed significant expression but were not detected as seasonality-induced genes.

| Tissue | Cluster | *PHT1* | *PHT5* | *PHO* |
| --- | --- | --- | --- | --- |
| The 6<sup>th</sup><br>leaf | A |  | Potri.014G078000 | *PHO12a* |
|  | B | *PHT1;11* | Potri.003G120600 | *PHO6, PHO11* |
|  | C | *PHT1;3, PHT1;7,*<br>*PHT1;8, PHT1;9* | Potri.001G111400 | *PHO3* |
|  | non-SI | *PHT1;4, PHT1;12* |  | *PHO4, PHO5,* |
| Stem | A | *PHT1;3* |  | *PHO3, PHO4, PHO6,*<br>*PHO11* |
|  | B |  | Potri.014G078000 | *PHO5, PHO7, PHO8,*<br>*PHO9, PHO12a* |
|  | C | *PHT1;4, PHT1;7* | Potri.003G120600 |  |
|  | D |  |  |  |
|  | E | *PHT1;9, PHT1;11,*<br>*PHT1;12* |  |  |
|  | non-SI |  | Potri.001G111400 |  |

In the 6^th^ leaf, none of *PHT1* family genes were classified in cluster A (Table 1, Fig. S6a). *PHT1;11* showed highest gene expression in stage 2, and was classified in cluster B. *PHT1;3*, *1;7*, *1;8* and *1;9* showed high gene expression in stage 3 (senescing season), and were classified in cluster C. Although *PHT1;4* and *1;12* were expressed at all sampling points, they were not detected as SI genes. Each of the three putative *PHT5* family genes showed different seasonal expression patterns and was classified into different clusters (Fig.7a, Table 1 and S6b). Potri.001G111400.1 showed high gene expression during stage 3, and was classified in cluster C. On the other hand, gene expression of Potri.003G120600.1 and Potri.014G078000.1 were high during stage 2, and were classified in cluster B and A, respectively.

Among *PHO* family genes which partly function for P outward transport from the cell, the expression of *PHO6*, *PHO11* and *PHO12a* decreased at 5^th^ week of stage3 (Fig. 7a and S7a). *PHO3* showed high gene expression during stage 3, and was classified in cluster C. *PHO4* and *PHO5* were expressed at all sampling points, but no seasonal oscillation was detected.

### Expression patterns of phosphate transporters in the stem

Among *PHT1* family genes, *PHT1;3* showed high gene expression during stage1 to 2, and was classified in cluster A (Fig. 7b, S8a and Table1). *PHT1;4* and *1;7* showed relatively high gene expression during the 3^rd^ to 8^th^ weeks of stage 3, and were classified in cluster C. However, since the seasonal oscillations of these genes (*PHT1;4* and *1;7*) were not clear, future studies will be needed to prove the existence of seasonal oscillations. *PHT1;9*, *1;11* showed high gene expression at Stage 1 on the second cycle. *PHT1;12* showed high gene expression at Stage 1 on both the first and the second cycle. These genes were classified in cluster E.

Among *PHT5*s (Fig.7b, S8b and Table1), Potri.001G111400.1 was expressed at all sampling points, but no seasonal oscillation was detected. Potri.003G120600.1 was classified in cluster C, and its expression decreased from stage 1 to the 3^rd^ week of stage 3, then started to increase and showed highest gene expression at the 8^th^ week of stage 3. Potri.014G078000.1 showed high gene expression during stage 1 to 2, and was classified in cluster B.

In *PHO* family genes, there were two major trends in expression patterns. (Fig. 7b, S9 and Table 1). *PHO3,4,6* and *11* were classified in cluster A, and showed highest gene expression at stage 1 and then decreased to dormancy stages. *PHO5*, *7-9* and *12a* were classified in cluster B, which showed high gene expression during stage 1 to the 3^rd^ week of stage 3 (Fig. 6b). Among them, gene expressions of *PHO7-9* increased stage 1 to stage 2.

In both the 6^th^ leaf and stem, several phosphate transporters showed a seasonal oscillating expression pattern in the shortened annual cycle system. However, since the gene expression of some genes (e.g., *PHT1;9* and *1;11* in stem) was relatively low, detailed quantification will be required in the future.

## Discussion

### Phosphorus re-translocation and overwintering recycling in the shortened annual cycle system

During the growing season, the 6^th^ leaf functioned as a source, and P was re-translocated to the apex and younger leaves mainly via phloem (Fig.3). Additionally, phloem-xylem exchange and xylem transport also occurred. Such circulation using both phloem and xylem to transport nutrients coincides with previous reports in herbaceous plants (Biddulph *et al*., 1958b) and reports about other nutrients in woody plants (Hartmann *et al*., 2000; Herschbach *et al*., 2012). Pfautsch *et al*. demonstrated by injection of a fluorescent dye that the radial transfer of water from phloem into xylem occurs predominantly via the symplast of ray parenchyma cells in *Eucalyptus saligna* (Pfautsch *et al*., 2015). Radial P transport may also occur symplastically along with the water flow. Phloem-xylem exchange would be an advantage especially for long-distance transport in tall trees. Because the xylem flow velocity (1.60 ± 0.09 mm s^-1^) is much faster than in the phloem (Windt *et al*., 2006), remobilized nutrients can be transported to upper parts in a shorter period.

In addition to apex and younger leaves, the stem cambium may also have sink ability because the stem diameter increased at stage 1 (Baba *et al*., 2018), and ^33^P was detected in the entire section including the cambium region (Fig. 4c).

During stage 1 to stage 3, seasonal shifts of sink tissues and of longitudinal re-translocation routes occurred. In the shortened annual cycle system, the apical meristem is active and trees keep growing in stage 1. Then, during stages 2 to 3, trees cease apical growth and leaves start to senesce. In the northern red oak, it was observed that the pattern of C allocation changed with the timing of flushing of new leaves (Dickson, 1989). Loss of sink ability in the apex and younger leaves may be one of the causes of the seasonal transition of re-translocation routes.

During the senescing season, the remobilized P was accumulated in phloem and xylem parenchyma cells including ray cells and perimedullary cells (Fig. 6). We have previously reported that winter P is stored as IP_6_ in the globoid-like structures in phloem and xylem parenchyma cells in poplar twigs both in nature and in the present laboratory system (Kurita *et al*., 2017). Netzer *et al*. also reported P accumulation in the cambial zone and wood ray parenchyma cells of *Populus × canescens* during dormancy (Netzer *et al*., 2017a). ^33^P localization using microautoradiography (Fig. 4d) agreed with these previous observations.

### Velocity of P re-translocation

Real-time imaging revealed a rapid flow of phosphate in poplar plants (Fig. 1). This rapid P re-translocation probably occurred via phloem (Fig. 3). The mechanism of phloem transport is generally explained by the Münch Theory as mass flow driven by an osmotically generated pressure gradient (Münch, 1930). Windt *et al*. measured velocities of the downward phloem flow in poplar using NMR imaging, and recorded a velocity of 0.34 ± 0.03 mm s^-1^ in the daytime (Windt *et al*., 2006). Schepper *et al*. found that phloem speed in angiosperms is roughly within the same range, around 0.17 mm s^-1^ (De Schepper *et al*., 2013). By the detection of ^32^P signals, we estimated that the transport velocity was about 0.015 mm s^-1^ to 0.021 mm s^-1^ (Fig. S2). This value is considerably slower than what has been reported and probably an underestimate. In poplar stems, ^32^P was localized in internal tissues (primarily the phloem), therefore its detection might be hindered in inverse proportion to the thickness of the outer tissues. It is believed that this decrease in signal intensity caused the signal to fall below the detection limit, leading to a delay in the detection of small amounts of ^32^P present in the tissue immediately after ^32^P administration. Therefore, our results only prove that P is transported at least as fast as this velocity. The measurement of the true velocity of P transport is a future challenge, and improvements in measurement techniques are expected.

### Seasonal expression pattern of phosphate transporters

Gene expression of phosphate (Pi) transporters showed seasonal oscillation (Fig. 7ab, S6-9), and their oscillations are thought to be related to the seasonal changes in P re-translocation route as observed by radioisotope imaging in this study (Fig. 1-5).

Through the entire stages, the 6^th^ leaf acted as a P source for young leaves or stem and roots (Fig. 2). Some Pi transporters showed constant expression, while others were highly expressed at a particular stage (Fig. 7a, Table1). These results suggest that even though a mature leaf consistently functions as a source, poplar probably optimizes seasonal P remobilization from leaves by using various Pi transporters in different combinations. Genes such as *PHT1;3 and 1;8*, which were classified in cluster C, may function to remobilize additional P from leaves before leaf fall. In a previous study, Loth-Pereda *et al*. reported that five *PHT1* members (*PHT1;1, 1;5, 1;6, 1;9* and *1;12)* were up-regulated in leaves of an adult poplar (*Populus trichocarpa*) during autumnal senescence (Loth-Pereda *et al*., 2011). These are not a perfect match to the *PHT1* members classified in cluster C. There are several possible causes for this difference (tree age, tree species, field or artificial experimental system, etc.). Further research is needed to reach more general conclusions, and here we only discuss the relationship to the P re-translocation observed in the experimental system.

In stem, Pi transporters also showed different expression patterns (Fig. 7b, S8 and S9). Although the velocity and direction of phloem flow itself is probably affected by photosynthates rather than P, Pi transporters are thought to be involved in seasonal P re-translocation by regulating the loading and unloading of P into sieve tubes and/or xylem vessels and the transport of P to surrounding parenchyma cells and vacuoles. *AtPHO1* is considered to be a Pi efflux transporter, and is responsible for xylem loading in roots (Hamburger *et al*., 2002; Arpat *et al*., 2012; Chiou, 2020). Recently, Che *et al*. reported that *OsPHO1;1* and *OsPHO1;2* showed influx Pi transporter activities, and taking their localization into account, *OsPHO1;1* and *PHO1;2* are thought to be involved in reloading P into the phloem and unloading from the xylem in the uppermost node connecting to the panicle, respectively (Che *et al*., 2020). *PtPHO*s classified into Cluster A and B may have the ability to seasonally regulate loading and unloading into and out of xylem vessels and sieve tubes.

In both the 6^th^ leaf and stem, each of the three members of *PtPHT5* showed different expression patterns. Potri.003G120600.1, which is markedly upregulated in stage 3, may contribute to uptake of intracellular P into the vacuole. Further studies to identify the localization and transport direction of each member will help to elucidate the role of *PtPHT5* in seasonal P re-translocation.

Besides the Pi transporters described here, SPDT members are reported to work as Pi transporter in rice and Arabidopsis (Ding *et al*., 2020). In poplar, these transporters may play an important role in P re-translocation. Our ongoing research will hopefully clarify their involvement.

## Conclusion

Seasonal P recycling is thought to be an adaptive strategy tailored to the lifestyle of deciduous plants. In this study, we successfully visualized the seasonal change of P re-translocation routes in poplar using a shortened annual cycle system, and clearly showed that re-translocated P from a leaf was reused for spring growth after overwintering. The expression profile of the Pi transporters was also obtained. These allow us to investigate the molecular mechanisms of seasonal P translocation in the same reproducible experimental system. Although the visualization of P re-translocation can only be conducted on small juvenile trees in a laboratory, the elucidation of the functions of Pi transporters and other components involved in seasonal P re-translocation will lead to an understanding of the annual phosphorus cycle in the large trees in field.

## Acknowledgement

We would like to offer our heartfelt thanks to Prof. Rob Reid (University of Adelaide, Australia) for his kind discussion and correction of this manuscript, Dr. Tatsuaki Goh and Dr. Koichi Toyokura for their kind discussion and incisive comments, Mr. Motohiro Mihara and Mr. Hitoshi Ooshima of Dynacom Co. Ltd. for their support in RNA-Seq analysis, Dr. Makoto Kashima, Dr. Mari Kamitani and Dr. Yoichi Hashida for their help in RNA-Seq analysis. Y. Kurita is deeply grateful to the SUNBOR Scholarship. This work was supported by a Grant-in-Aid for Scientific Research of Innovative Areas from the Japanese Ministry of Education, Sports, Culture, Science, and Technology on ‘‘Perceptive plants (22120006)’’ and “Plant-structure-opt (18H05493)” for TM, JSPS KAKENHI grant Number 15J03992 and 18K14735 for YK, and JST PREST grant Number JPMJPR15Q7 for KT. The authors declare no competing financial interests.

## Author Contribution

YK, TM and AN designed the study and wrote the manuscript. YK performed all RI experiments and RNA-Seq analysis under the instruction of following co-authors. SK and RS instructed and performed experiments using the real-time radioisotope imaging system. AH instructed and performed experiments using MAR. KT and TMN advised experimental design using RRIS and MAR. AT, AD and YK prepared RNA-Seq library. AN instructed RNA-Seq analysis. KB instructed on dendrology. MO, HF and KI supported YK’s experiments in Kobe University. All authors discussed the results and advised on the manuscript.

## Data Availability

Sequence data of RNA-Seq have been submitted to the NCBI Sequence Read Archive repository under the BioProject number, PRJNA756807 (https://www.ncbi.nlm.nih.gov/bioproject/756807). R scripts and data required for analysis (count data, etc.) are available via the GitHub repository (https://github.com/YukoKurita/2021_RI_SACsystem_RNA-seq_Analysis).

## Figure legends of Supporting Information

**Fig. S1** Surgical treatments on *Populus alba* trees and a summary diagram of culture conditions.

**Fig. S2** Transition of ^32^P signal intensity (intensity cm^-2^) of the apex and the 6^th^ leaf axil.

**Fig. S3** ^33^P autoradiographs of leaf axil of the 6^th^ leaf in stage 1 using imaging plate.

**Fig. S4** Microautoradiographs of the stem between 5^th^ and 6^th^ nodes.

**Fig. S5** Summary of sample sets and gene sets in this study.

**Fig. S6** Expression plots of *PHT* family genes in the 6^th^ leaf.

**Fig. S7** Expression plots of *PHO* family genes and typical seasonally variable genes in the 6^th^ leaf.

**Fig. S8** Expression plots of *PHT* family genes in the stem.

**Fig. S9** Expression plots of *PHO* family genes in the stem.

**Video S1** Real-time imaging movie of ^32^P in *Populus alba* at stage 1.

**Video S2** Real-time imaging movie of ^32^P in *Populus alba* at stage 3.

**Video S3** 3D image of the 6^th^ axil of *Populus alba* at stage 1.

**Video S4** 3D image of the 6^th^ axil of *Populus alba* at stage 3.

