## Supporting Information for "Visualization of seasonal phosphorus re-translocation, and expression profile of phosphate transporters in a shortened annual cycle system of the deciduous poplar tree"

The following Supporting Information is available for this article:

**Fig. S1** Surgical treatments on *Populus alba* trees and a summary diagram of culture conditions.

**Fig. S2** Transition of ^32^P signal intensity (intensity cm^-2^) of the apex and the 6^th^ leaf axil.

**Fig. S3** ^33^P autoradiographs of leaf axil of the 6^th^ leaf in stage 1 using imaging plate.

**Fig. S4** Microautoradiographs of the stem between 5^th^ and 6^th^ nodes.

**Fig. S5** Summary of sample sets and gene sets in this study.

**Fig. S6** Expression plots of *PHT* family genes in the 6^th^ leaf.

**Fig. S7** Expression plots of *PHO* family genes and typical seasonally variable genes in the 6^th^ leaf.

**Fig. S8** Expression plots of *PHT* family genes in the stem.

**Fig. S9** Expression plots of *PHO* family genes in the stem.

**Video S1** Real-time imaging movie of ^32^P in *Populus alba* at stage 1.

**Video S2** Real-time imaging movie of ^32^P in *Populus alba* at stage 3.

**Video S3** 3D image of the 6^th^ axil of *Populus alba* at stage 1.

**Video S4** 3D image of the 6^th^ axil of *Populus alba* at stage 3.

The following Supporting Information Tables are available as a single Excel file, submitted separately:

**Table S1** Attributes of the samples used in the present study

**Table S2**  List of expressed genes and seasonality-induced genes in the 6^th^ leaf.

**Table S3** List of expressed genes and seasonality-induced genes in stem.

**Table S4** The result of GO enrichment analysis of seasonality induced genes in the 6^th^ leaf classified into cluster A.

**Table S5** The result of GO enrichment analysis of seasonality induced genes in the 6^th^ leaf classified into cluster B.

**Table S6** The result of GO enrichment analysis of seasonality induced genes in the 6^th^ leaf classified into cluster C.

**Table S7** The result of GO enrichment analysis of seasonality induced genes in stem classified into cluster A.

**Table S8** The result of GO enrichment analysis of seasonality induced genes in stem classified into cluster B.

**Table S9**  The result of GO enrichment analysis of seasonality induced genes in stem classified into cluster C.

**Table S10** The result of GO enrichment analysis of seasonality induced genes in stem classified into cluster D.

**Table S11** The result of GO enrichment analysis of seasonality induced genes in stem classified into cluster E.

**Fig. S1** Surgical treatments on *Populus alba* trees and a summary diagram of culture conditions. (a) How to apply ^32^P or ^33^P to the 6^th^ leaf. A ‘flap’ was cut along a vein from the basal direction. The flap was covered with a pipette tip containing ^32^P or ^33^P solution. (b) Enlarged view of phloem girdling. Orange arrows indicate the position of phloem girdling. (c) Summary diagram of culture conditions in the shortened annual cycle system.

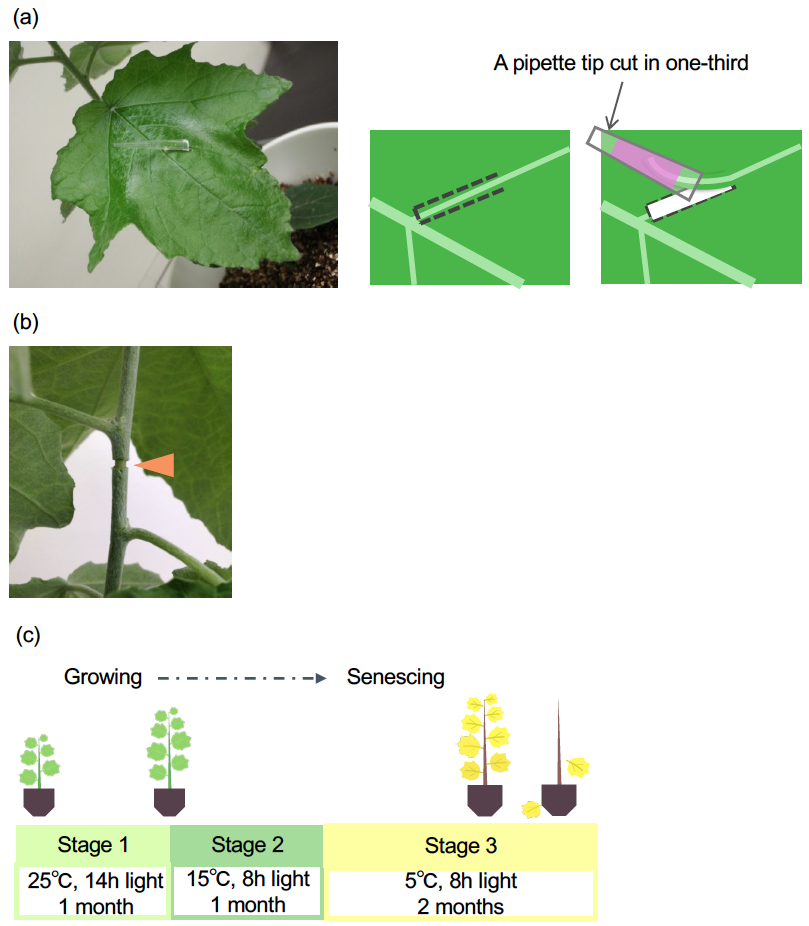

**Fig. S2** Transition of ^32^P signal intensity (intensity / cm^2^) of the apex and the 6^th^ leaf axil. (a)(b) Time course plot of ^32^P signal in tree A and tree B. Pink lines and blue lines indicate the trend line of signal intensity at the 6^th^ leaf axil and apex, respectively. Trend lines were drawn using the smooth.spline function of R. Black broken lines indicate the detection threshold. (c)(d) The ROIs of measurement in tree A and tree B, respectively. 1: Apex, 2: The 6^th^ leaf axil. 3-5: Background.

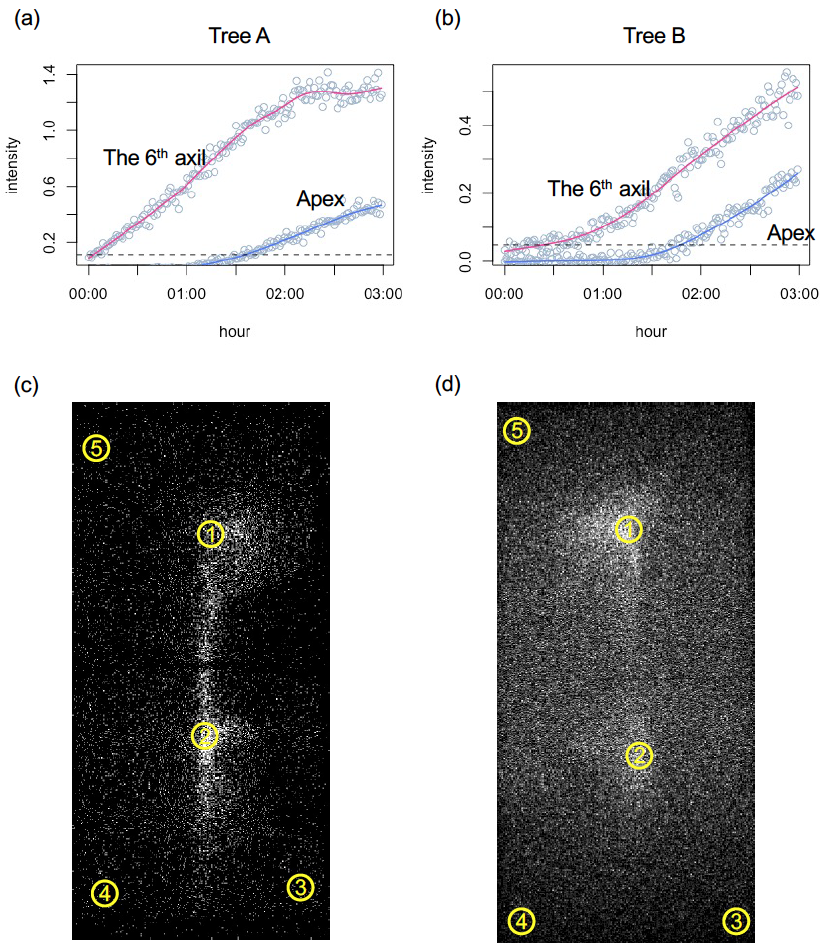

**Fig. S3** ^33^P autoradiographs of leaf axil of the 6^th^ leaf in stage 1 using imaging plate. (a) Autoradiographs of leaf axil of the 6^th^ leaf in stage 1. (b) Autoradiographs of leaf axil of the 6^th^ leaf in stage 3. From left to right, sections from upper to lower part of the plant.

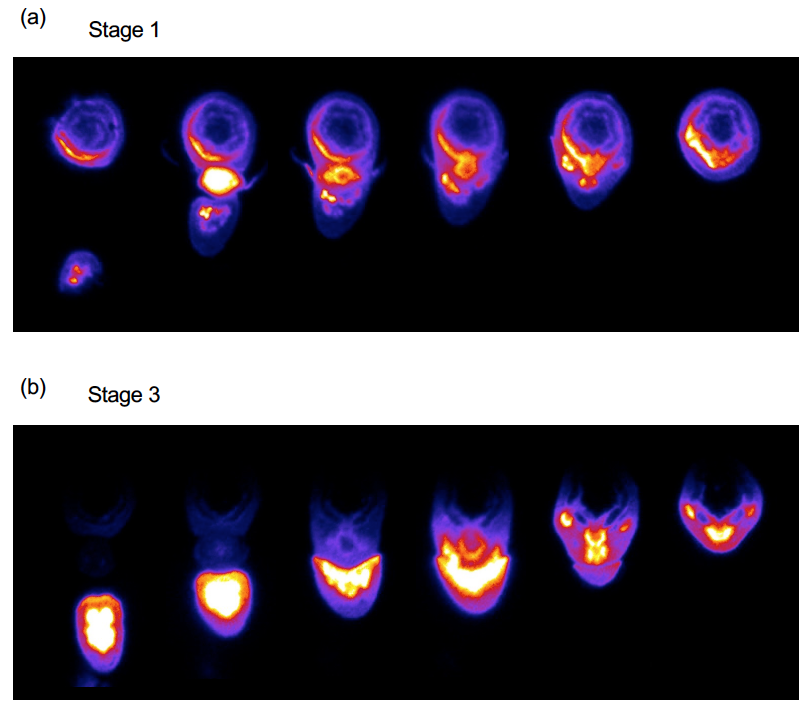

**Fig. S4** Microautoradiographs of the stem between 5^th^ and 6^th^ nodes. (a) The stem between 5^th^ and 6^th^ nodes in stage 1. (b) The stem between 9^th^ and 10^th^ nodes in stage 1. (c) The stem between 5^th^ and 6^th^ nodes in stage 3. (d) The stem between 9^th^ and 10^th^ nodes in stage 3. Left: the section stained by toluidine blue. Middle: microautoradiograph of the section. Right: merged image. Green color arrowheads indicate the position of the 6th leaf. Bar = 0.5 mm.

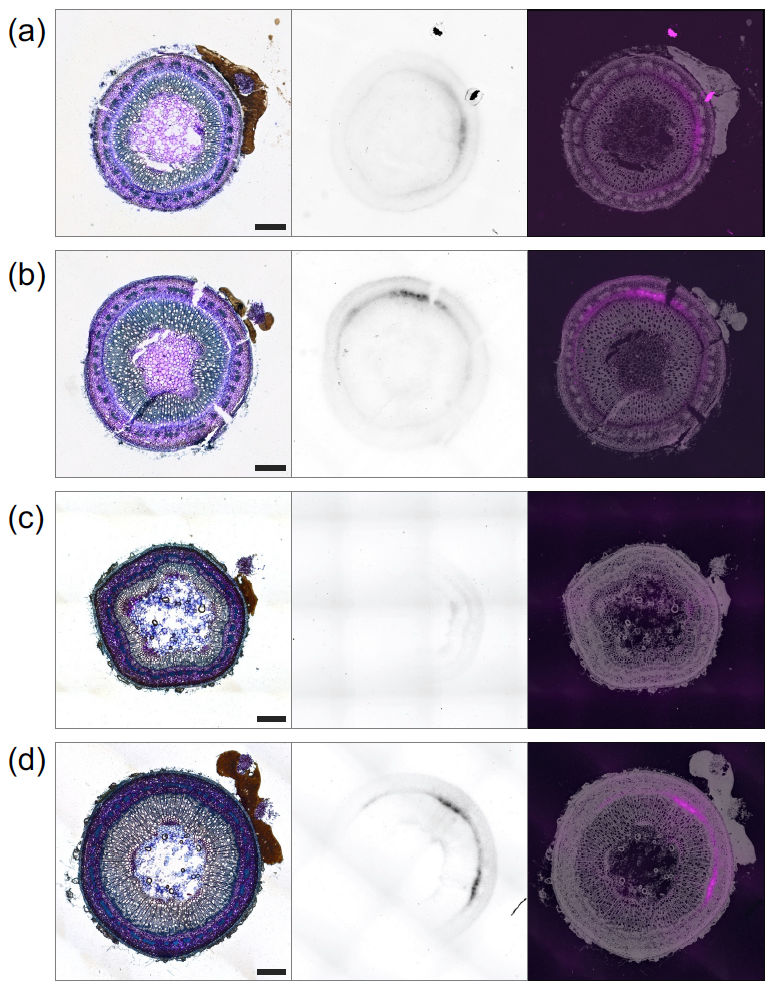

**Fig. S5** Summary of sample sets and gene sets in this study. (a) Sample sets of this study. (b) Summary of gene sets in this study. The number of genes written in green on the left and in orange on the right are the numbers of genes defined in the 6^th^ leaf and stem sample sets, respectively.

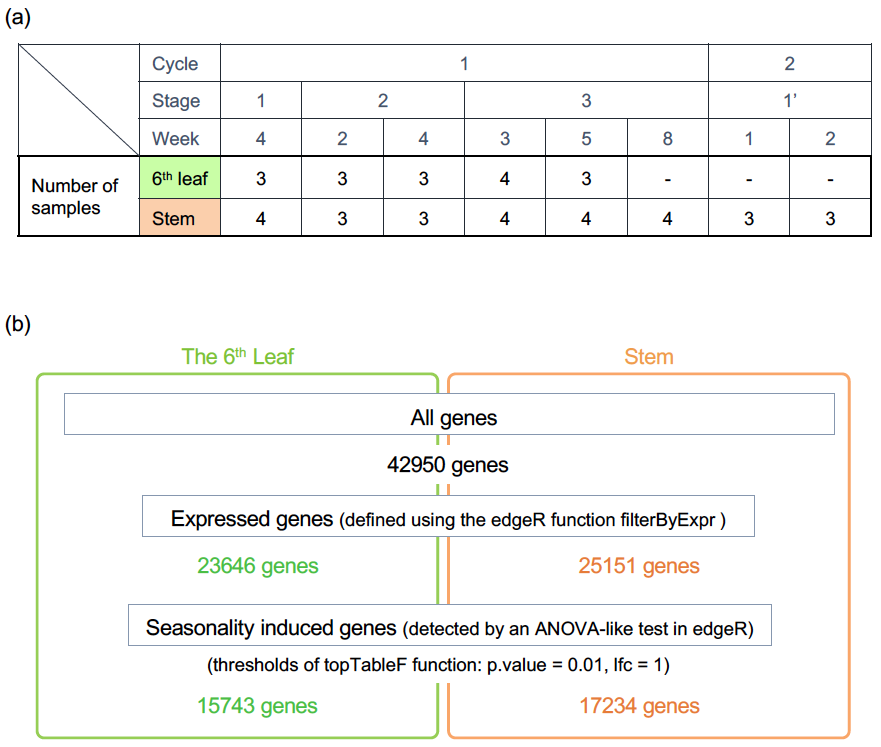

**Fig. S6** Expression plots of *PHT* family genes in the 6^th^ leaf. (a) *PHT1* family genes which were detected as the expressed genes. For genes detected as seasonality-induced genes, the names of the clusters are shown in the upper left corner of the graph. (b) Homolog genes of *AtPHT5* genes which were detected as the expressed genes. The names of the *AtPHT5* genes with the highest similarity in the Photozome database are listed next to the transcript name *P. trichocarpa*. Different letters in plots indicate differences between sampling points at *P* < 0.05 using ANOVA and Tukey’s HSD test.

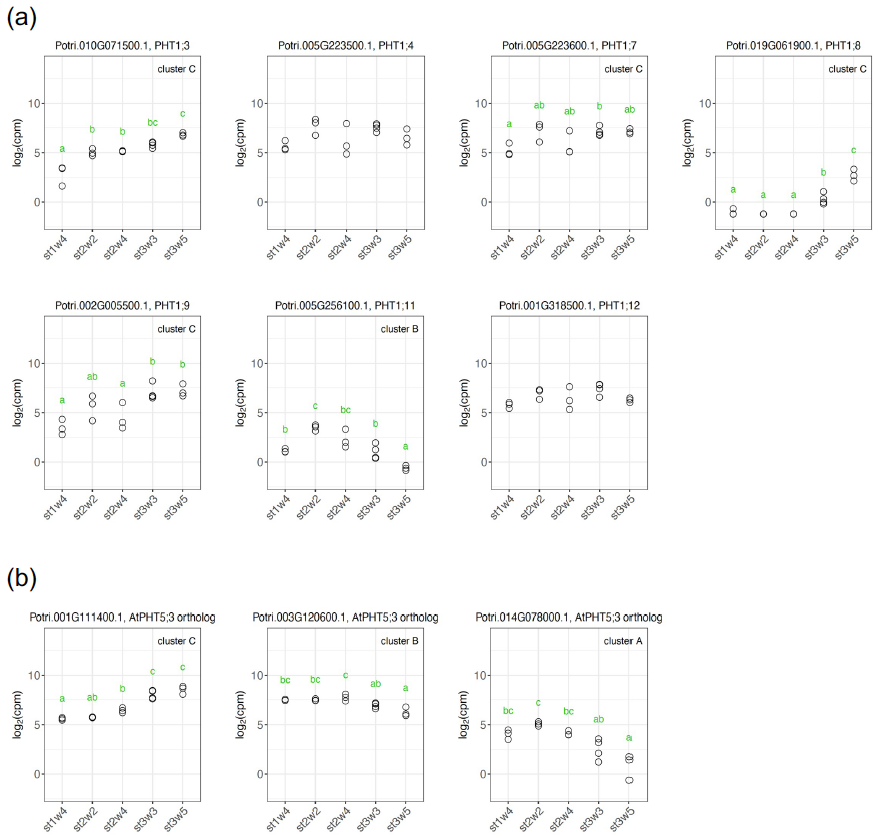

**Fig. S7** Expression plots of *PHO* family genes and typical seasonally variable genes in the 6^th^ leaf. (a) *PHO* family genes which were detected as expressed genes. For genes detected as seasonality-induced genes, the names of the clusters are shown in the upper left corner of the graph. *PHO12* which was annotated by Zhang et al. (2016) was split into two transcripts, Potri.T048933.1 and Potri.T097866.1, due to a database update. Therefore, we analyzed the expression of these two transcripts (named as *PHO12a* and *PHO12b*, respectively). (b) Expression plots of genes whose seasonal expression patterns have been clarified in previous studies. Different letters in plots indicate differences between sampling points at *P* < 0.05 using ANOVA and Tukey’s HSD test.

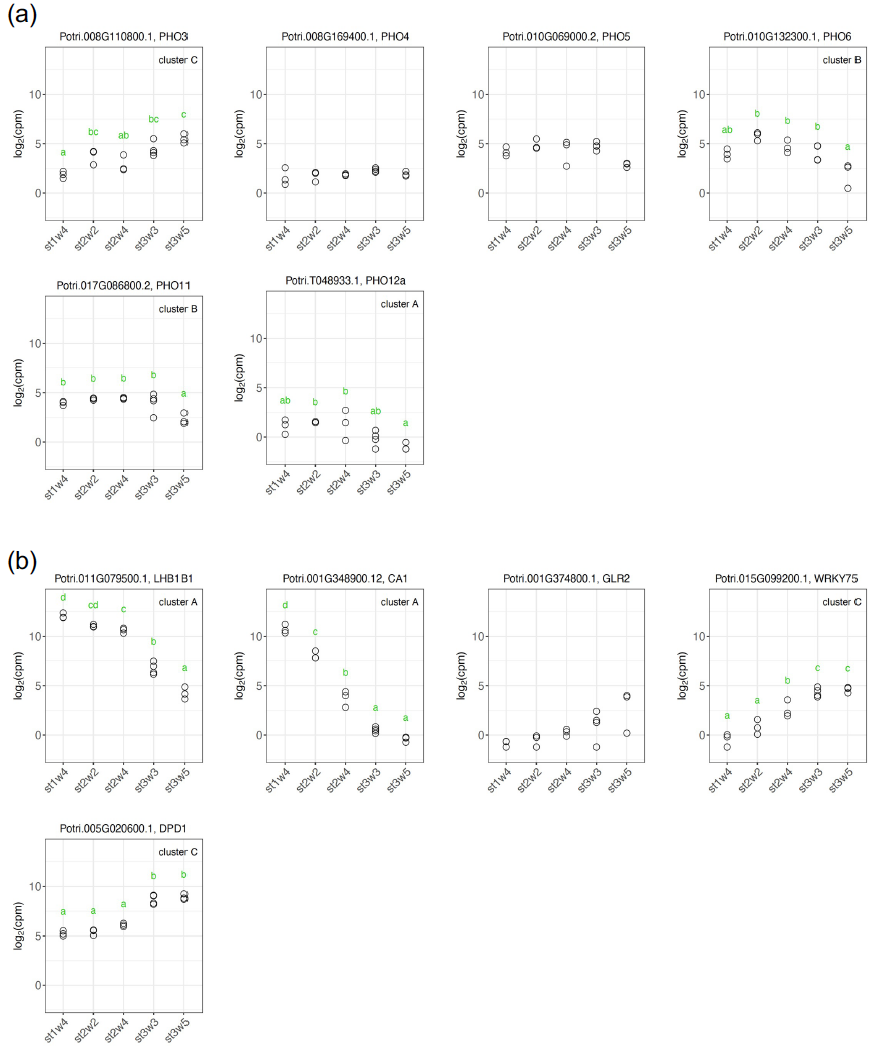

**Fig. S8** Expression plots of *PHT* family genes in the stem. (a) *PHT1* family genes which were detected as expressed genes. For genes detected as seasonality-induced genes, the names of the clusters are shown in the upper left corner of the graph. (b) Homolog genes of *AtPHT5* genes which were detected as expressed genes. The names of the *AtPHT5* genes with the highest similarity in the Photozome database are listed next to the transcript name of *P. trichocarpa*. Different letters in plots indicate differences between sampling points at *P* < 0.05 using ANOVA and Tukey’s HSD test.

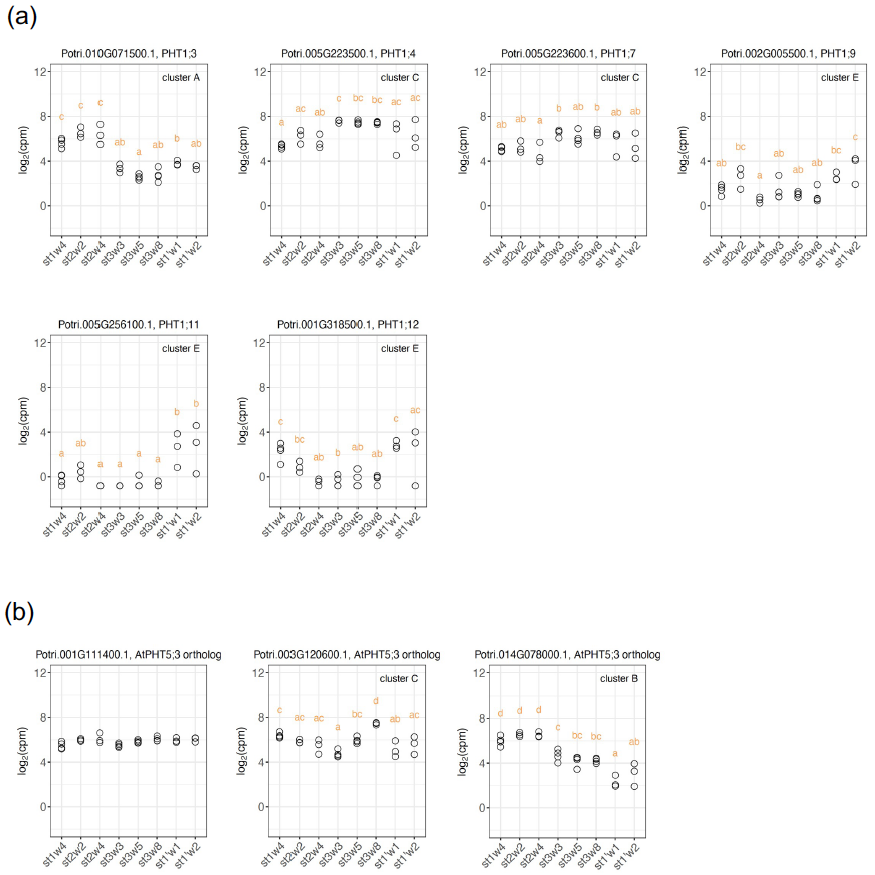

**Fig. S9** Expression plots of *PHO* family genes in the stem which were detected as expressed genes. For genes detected as seasonality-induced genes, the names of the clusters are shown in the upper left corner of the graph. Different letters in plots indicate differences between sampling points at *P* < 0.05 using ANOVA and Tukey’s HSD test.

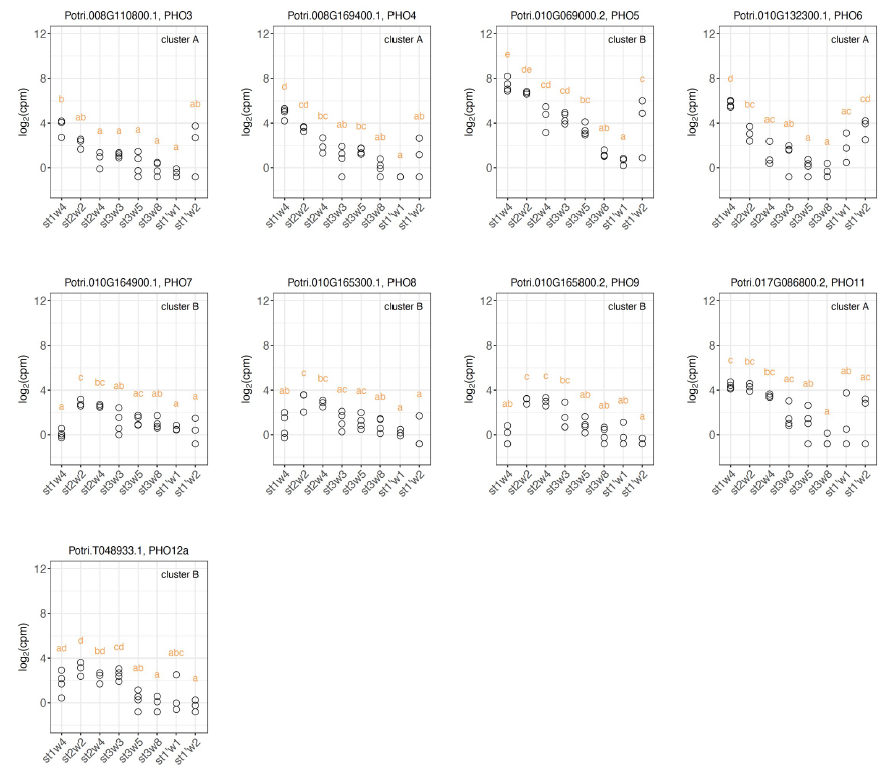
